# Long-read sequencing resolves structural variants in *SERPINC1* causing antithrombin deficiency and identifies a complex rearrangement and a retrotransposon insertion not characterized by routine diagnostic methods

**DOI:** 10.1101/2020.08.28.271932

**Authors:** Belén de la Morena-Barrio, Jonathan Stephens, María Eugenia de la Morena-Barrio, Luca Stefanucci, José Padilla, Antonia Miñano, Nicholas Gleadall, Juan Luis García, María Fernanda López-Fernández, Pierre-Emmanuel Morange, Marja K Puurunen, Anetta Undas, Francisco Vidal, NIHR BioResource, F Lucy Raymond, Vicente Vicente García, Willem H Ouwehand, Javier Corral, Alba Sanchis-Juan

## Abstract

The identification and characterization of structural variants (SVs) in clinical genetics have remained historically challenging as routine genetic diagnostic techniques have limited ability to evaluate repetitive regions and SVs. Long-read whole-genome sequencing (LR-WGS) has emerged as a powerful approach to resolve SVs. Here, we used LR-WGS to study 19 unrelated cases with type I Antithrombin Deficiency (ATD), the most severe thrombophilia, where routine molecular tests were either negative, ambiguous, or not fully characterized. We developed an analysis workflow to identify disease-associated SVs and resolved 10 cases. For the first time, we identified a germline complex rearrangement involved in ATD previously misclassified as a deletion. Additionally, we provided molecular diagnoses for two unresolved individuals that harbored a novel SINE-VNTR-Alu retroelement insertion that we fully characterized by *de novo* assembly and confirmed by PCR amplification in all affected relatives. Finally, the nucleotide-level resolution achieved for all the SVs allowed breakpoint analysis, which revealed a replication-based mechanism for most of the cases. Our study underscores the utility of LR-WGS as a complementary diagnostic method to identify, characterize, and unveil the molecular mechanism of formation of disease-causing SVs, and facilitates decision making about long-term thromboprophylaxis in ATD patients.

## Main text

Haploinsufficiency of *SERPINC1* (MIM: 107300) is associated with type I antithrombin deficiency (ATD), that constitutes the most severe thrombophilia since it significantly increases the risk of venous thrombosis (OR:20-30).^2^ Routine investigation of ATD combines functional assays, antigen quantification and genetic analyses. Causal variants are identified in *SERPINC1* for 70% of cases, whilst 5% of patients harbor defects in other genes and 25% remain without a genetic diagnosis.^2^ The majority of reported pathogenic variants in *SERPINC1* are small genetic defects (63% single-nucleotide variants and 28% indels), with structural variants (SVs) accounting for a smaller proportion accounting for a smaller proportion of cases.^3; 4^

Structural variants are genomic rearrangements involving more than 50 nucleotides that contribute to genomic diversity and function, evolution, and can cause somatic and germline diseases.^5-7^ Despite improvements in genomic technologies, characterization of SVs remains challenging and the full spectrum of SVs is not achieved by routine methods such as microarrays or other targeted sequencing approaches. In ATD, the detection and characterization of SVs remain particularly challenging due to the high number of repetitive elements in and around *SERPINC1* (35% of *SERPINC1* sequence are interspersed repeats).^8^ Copy number variants causing ATD are routinely identified in specialized centers by multiplex ligation-dependent probe amplification (MLPA),^2^ but this technology does not consider the full spectrum of SVs. Additionally, it does not provide a nucleotide-level resolution, which is important for confirming causality and reveal insights into SVs formation.^9-11^ These limitations may now be addressed by long reads, that can span repetitive or other problematic regions, allowing identification and characterization of SVs.^10; 12-16^

Here, we report on the results of long-read whole-genome sequencing (LR-WGS) on 19 unrelated cases with ATD, where routine molecular tests were either negative, ambiguous, or did not fully characterize a SV, in order to identify, resolve and investigate the most likely molecular mechanism of formation of causal SVs involved in this severe thrombophilia.

Nineteen unrelated individuals with ATD were selected from our cohort of 340 cases, recruited between 1994 and 2019: 8 patients with causal SVs identified by MLPA were included for variant characterization and investigation of the potential mechanisms of formation, and 11 patients were selected because multiple independent genetic studies evaluating *SERPINC1* had failed to identify causal variants (Table S1, Supplementary Methods). Measurement of antithrombin levels and function were performed for all participants as previously described.^17; 18^ LR-WGS was performed using the PromethION platform (Oxford Nanopore Technologies) and a multi-modal analysis workflow for the sensitive detection of SVs was developed (http://github.com/who-blackbird/magpie) and applied (Figure 1A, Supplemental Methods). Detailed information is provided in Supplemental Methods.

**Figure 1.**
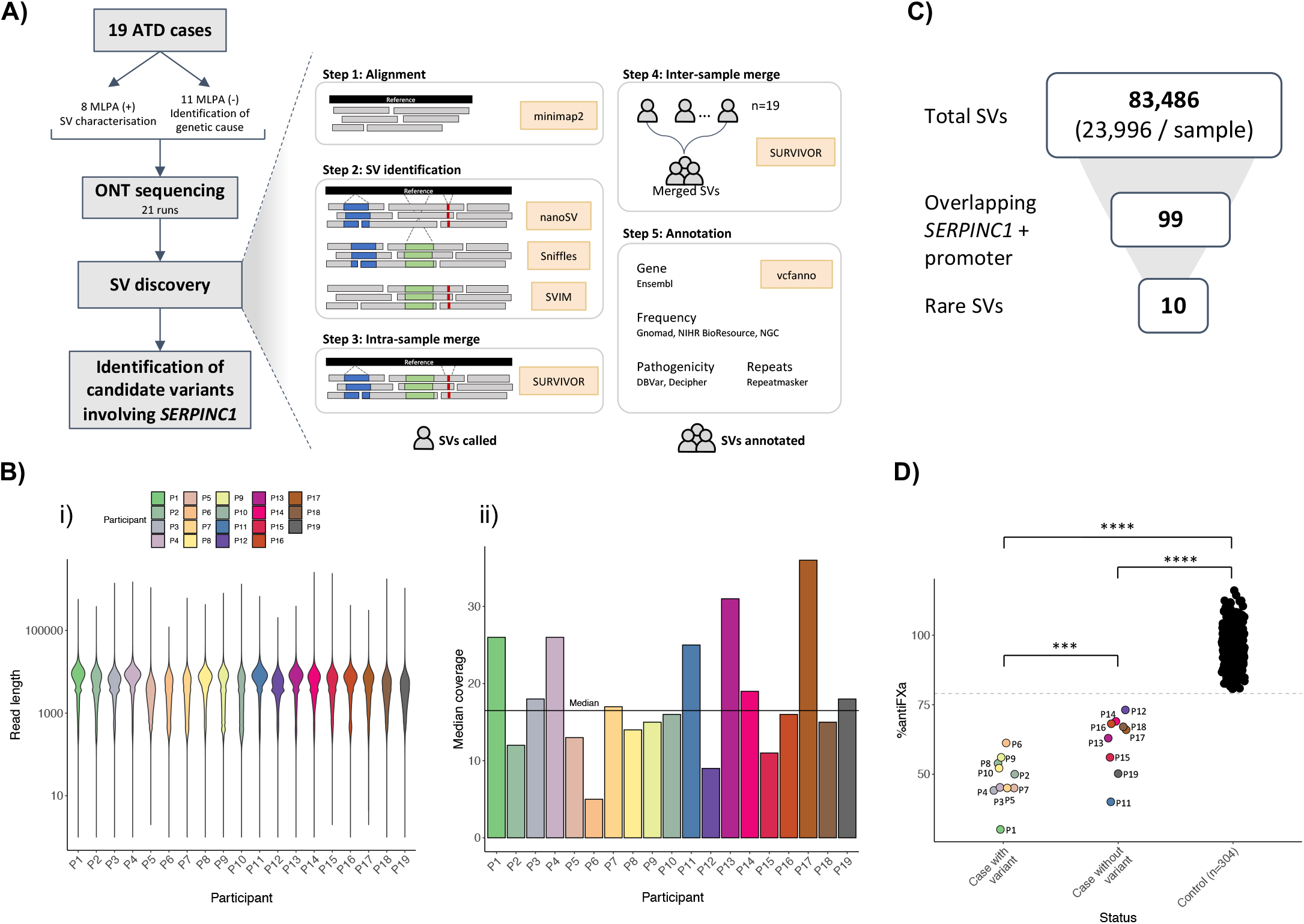
Long-read sequencing workflow and results. **(A)**Overview of the general stages of the SVs discovery workflow. Algorithms used are depicted in yellow boxes. **(B)** Nanopore sequencing results. i) Sequence length template distribution. Average read length was 4,499 bp (sd ± 4,268); the maximum read length observed was 2.5Mb. ii) Genome median coverage per participant. The average across all samples was 16x (sd ± 7.7). **(C)** Filtering approach and number of SVs obtained per step. SERPINC1 + promoter region corresponds to [GRCh38/hg38] Chr1:173,903,500-173,931,500. **(D)** anti-FXa percentage levels for the participants with a variant identified (P1-P10), cases without a candidate variant (P11-P19) and 300 controls from our internal database. The statistical significance is denoted by asterisks (*), where ***P<0.001, ****P≤ 0.0001. p-values calculated by one-way ANOVA with Tukey’s post-hoc test for repeated measures. ATD=Antithrombin Deficiency; ONT=Oxford Nanopore Technologies; SV=Structural Variant.

Nanopore sequencing in 21 runs produced reads with an average length of 4,499 bp and a median genome coverage of 16x (Figure 1A-B). After a detailed quality control analysis (Figure S1) 83,486 SVs were identified, consistent with previous reports using LR-WGS (Figure S2).^12^ Focusing on rare variants (allele count <= 10 in gnomAD v3, NIHR BioResource and NGC project) ^15; 19; 20^ in *SERPINC1* and flanking regions, 10 candidate heterozygous SVs were observed in 9 individuals (Figure 1C). Visual inspection of read alignments identified an additional heterozygous SV in a region of low coverage involving *SERPINC1*.

First, Nanopore sequencing resolved the precise configuration of SVs previously identified by MLPA in 8 individuals (P1-P8). SVs were identified independently of their size (from 7Kb to 968 Kb, restricted to *SERPINC1* or involving neighboring genes) and their type (six deletions, one tandem duplication and one complex SV) (Figure 2, Table S1). Importantly, SVs for two cases with previous inconsistent or ambiguous results were characterized by Nanopore sequencing (P2 and P6) (Figure 2, Table S1).

**Figure 2.**
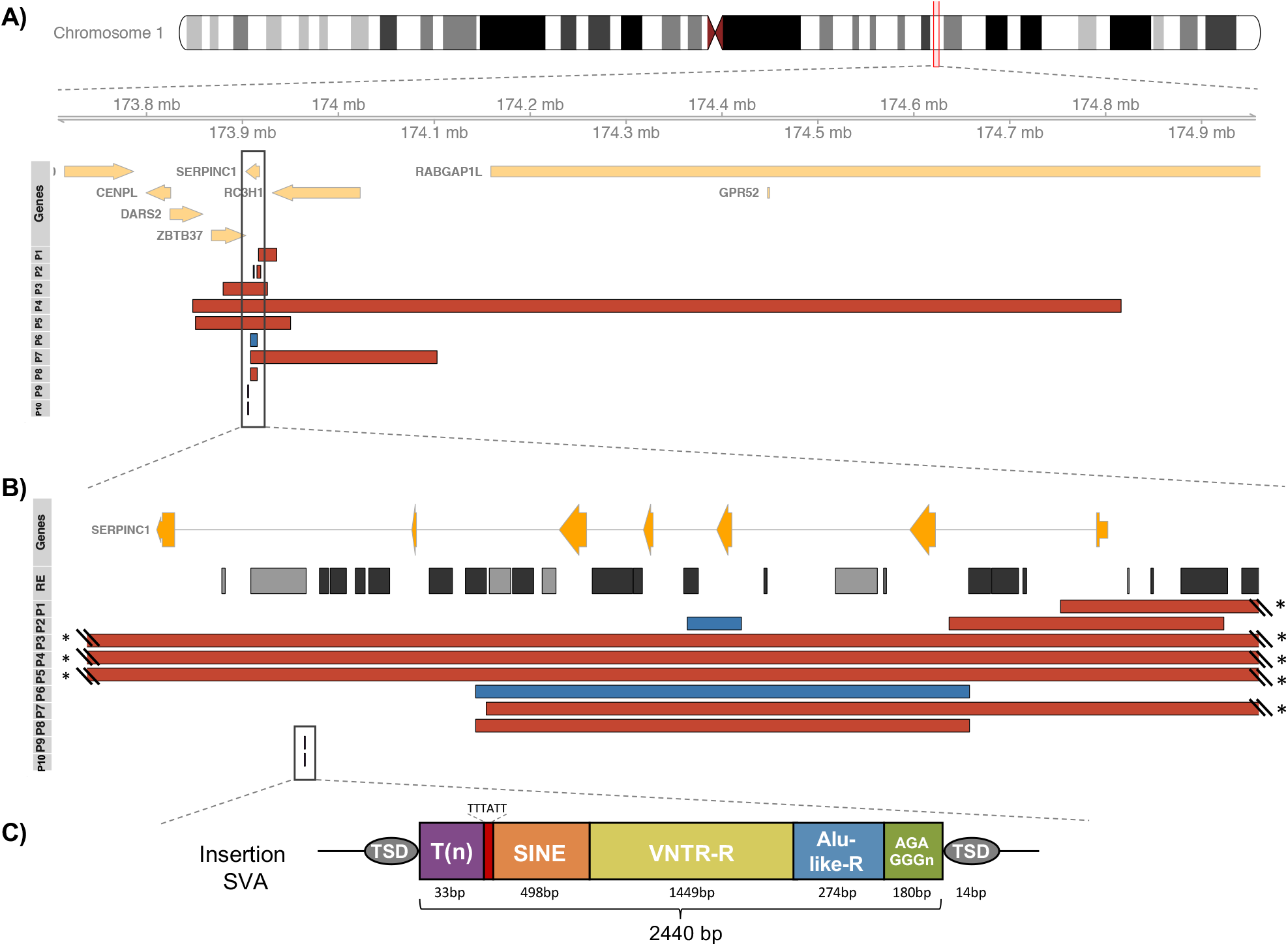
Candidate SVs identified by long-read sequencing. **(A)** Schematic of chromosome 1 followed by protein coding genes falling in the zoomed region (1q25.1). SVs for each participant (P) are colored in red (deletions) and blue (duplications). The insertion identified in P9 and P10 is shown with a black line. **(B)** Schematic of SERPINC1 gene (NM_000488) followed by repetitive elements (RE) in the region. SINEs and LINEs are colored in light and dark grey respectively. Asterisks are present where the corresponding breakpoint falls within a RE. **(C)** Characteristics of the antisense-oriented SVA retroelement (respect to the canonical sequence)^1^ observed in P9. Length of the fragments are subject to errors from nanopore sequencing. TSD=Target site duplication.

For the first case (P2), MLPA detected a deletion of exon 1, but long-range PCR followed by NGS suggested a deletion of exons 1 and 2. The discordant results were explained by a complex SV in *SERPINC1* revealed by Nanopore sequencing, that resulted in a dispersed duplication of exon 3 and the deletion of exon 1, both in the same allele (Figure S3A). Although complex SVs have already been associated with human disease,^10; 21^ this is the first report of a germline complex rearrangement involved in ATD, that was also confirmed by Sanger sequencing in the affected daughter of P2. Further investigations would be required to elucidate whether the complex SV was formed by one or two independent mutational events.

For the second case (P6), MLPA detected a duplication of exons 1, 2 and 4 and a deletion of exon6. Here, our sequencing approach identified a tandem duplication of exons 1 to 5, which was confirmed by long-range PCR (Figure S3B) and observed to be present in the affected son of P6.

Then, we aimed to identify new disease-causing variants in the remaining 11 participants. Remarkably, two cases (P9 and P10) presented an insertion of a SINE-VNTR-Alu (SVA) retroelement of 2,440 bp (Figure 2, Table S1), suspected to induce transcriptional interference of *SERPINC1. De novo* assembly using the sequencing data of P9 revealed an antisense-oriented SVA element flanked by a target site duplication (TSD) of 14 bp (Figure 2C), consistent with a target-primed reverse transcription mechanism of insertion into the genome.^1;22^ Interestingly, the TSD in both individuals was the same, suggesting a shared mechanism of formation or a founder effect. The inserted sequence was aligned to the canonical SVA A-F sequences (Figure S4A) and it was observed to be closest to the SVA E in the phylogenetic tree (Figure S4B). Moreover, the VNTR sub-element harbored 1,449bp, which was longer than the typical ∼520bp-long VNTR in the canonical sequences.

These results highlight the heterogeneous genomic landscape of SVA sequences and underscore the importance of their characterization in order to obtain a reliable catalogue of novel mobile elements to identify and interpret this type of causal variants in other patients and other disorders where retrotransposon insertions might also be involved.^1;23;24^ This characterization has been historically challenging by the application of classic technologies, but here we show that it can be achieved by *de novo* assembly of long-reads. The SVA insertion was confirmed in P9, P10 and two other affected relatives by specifically designed PCR amplification and Sanger sequencing facilitated by the Nanopore data (Figure S5). SVA retroelements are challenging to amplify given their genomic characteristics (GC-rich sequences and length). Here, multiple PCRs were attempted, and the final amplified product was only obtained by using an internal SVA primer (Figure S5).

Finally, breakpoint analysis was performed to investigate the mechanism underlying the formation of these SVs involving *SERPINC1*. Nanopore sequencing facilitated primer design to perform Sanger sequencing confirmations for all the new formed junctions, demonstrating a 100% accuracy in 7/10 (70%) SVs called. Repetitive elements (RE) were detected in all the SVs, with Alu elements being the most frequent (16/24, 67%) (Table S2). Alu-mediated SVs have been previously reported as associated with ATD, ^25^ and their frequency is consistent with the high proportion of Alu sequences in *SERPINC1* (22% of intronic sequence).^26^ Additionally, breakpoint analysis identified microhomologies (7/11, 64%) and insertions, deletions or duplications (7/11, 64%) (Figure S6).

Specific mutational signatures can yield insights into the mechanisms by which the SVs are formed. Our results suggest a replication-based mechanism (such as BIR/MMBIR/FoSTeS) for most of the cases (P1-P8).^27^ Importantly, we observed a non-random formation driven by the presence of REs in some of the SVs. For example, an *Alu* element in intron 1 was involved in the SVs of P6 and P8, and an *Alu* element in intron 5 was involved in SVs of P6, P7 and P8 (Figure 2B, Table S3). It has been suggested that RE may provide larger tracks of microhomologies, also termed ‘microhomology islands’, that could assist strand transfer or stimulate template switching during repair by a replication-based mechanism.^27^ These microhomology islands were present in the SVs of 4 cases (P4, P6-P8), highlighting the important role that RE play in the formation of non-recurrent, but non-random, SVs.

Overall, we resolved SVs affecting *SERPINC1* in 10 individuals with ATD. However, 9 additional cases remain as yet unresolved, three of whom reported to have familial disease. An explanation may be that the causal variant was missed due to low coverage, or alternatively the variant is located in an unidentified transacting gene or in a regulatory element for *SERPINC1*, as we have recently reported for other genes.^15^ The observation that the ATD patients without causal SVs have significantly higher anti-FXa activity than those with SVs (Figure 1D) is supportive of the notion that causal variants may regulate gene expression.

Here, we show how LR-WGS can be used to resolve SVs causal of ATD, independently of the length or the type, that can be missed, misunderstood or misclassified by routine molecular diagnostic methods. Moreover, we report for the first time a germline complex rearrangement and the insertion of a SVA retroelement as the genetic defect responsible of ATD and reveal insights into the mechanisms of formation of these SVs. Altogether this study highlights the importance of identifying a new class of causal variants to improve diagnostic rates, to provide accurate family counselling and to facilitate decision making about long-term thromboprophylaxis.

## Supplemental data

Supplemental methods, figures and tables are provided in separate documents, which will be linked directly from *bioRxiv*.

## Acknowledgments

We thank the participants involved in this study and their families. We thank NIHR BioResource volunteers for their participation, and gratefully acknowledge NIHR BioResource centers, NHS Trusts and staff for their contribution. We thank the National Institute for Health Research, NHS Blood and Transplant, and Health Data Research UK as part of the Digital Innovation Hub Programme. The views expressed are those of the author(s) and not necessarily those of the NHS, the NIHR or the Department of Health and Social Care. This work was supported by the National Institute for Health Research England (NIHR) for the NIHR BioResource project (grant numbers RG65966 and RG94028), the PI18/00598 project (ISCIII y FEDER) and the 19873/GERM/15 project (Fundación Séneca). All participants provided written informed consent to participate in the study. The study was approved by the East of England Cambridge South national institutional review board (13/EE/0325). The research conforms to the principles of the Declaration of Helsinki.

## Data and Code Availability

Sequence data for all the individuals in this work have been deposited at the European Genome-Phenome Archive under the accession number EGAD00001006254. The workflow developed for the detection of SVs is publicly available at http://github.com/who-blackbird/magpie.

## Authorship Contributions

BMB, WHO, JC and ASJ designed the study. MMB, LS, JP, AM, NG, FLR and VV helped with study design. BMB, MMB, JP, AM performed laboratory experiments and analyzed the experimental data. JS performed sample preparation and executed long-read sequencing. ASJ developed the analysis workflow for long-read sequencing, applied this to data processing and performed the computational and statistical analyses. BMB performed computational analyses and variant validation. JM, FV, provided valuable insight into CGHa and NGS data analysis. AU, MF, MP and PM recruited participants and collected the clinical data and samples. BMB, WHO, JC and ASJ wrote the manuscript. All authors read and approved the final manuscript.

