## Supplementary Material for "Long-read sequencing resolves structural variants in *SERPINC1* causing antithrombin deficiency and identifies a complex rearrangement and a retrotransposon insertion not characterized by routine diagnostic methods"

##### Authors and Affiliations

Belén de la Morena-Barrio [1], Jonathan Stephens [2,3], María Eugenia de la Morena-Barrio [1], Luca Stefanucci [2,4,5], José Padilla [1], Antonia Miñano [1], Nicholas Gleadall [2,3], Juan Luis García [6], María Fernanda López-Fernández [7], Pierre-Emmanuel Morange [8], Marja K Puurunen [9], Anetta Undas [10], Francisco Vidal [11], NIHR BioResource [3], F Lucy Raymond [3,12], Vicente Vicente García [1], Willem H Ouwehand [2,3], Javier Corral [1,13], Alba Sanchis-Juan [2,3,13]

1. Servicio de Hematología y Oncología Médica, Hospital Universitario Morales Meseguer, Centro Regional de Hemodonación, Universidad de Murcia, Instituto Murciano de Investigación Biosanitaria (IMIB-Arrixaca), Centro de Investigación Biomédica en Red de Enfermedades Raras (CIBERER), Murcia, Spain.
2. Department of Haematology, University of Cambridge, NHS Blood and Transplant Centre, Cambridge, CB2 0PT, UK
3. NIHR BioResource, Cambridge University Hospitals NHS Foundation Trust, Cambridge Biomedical Campus, Cambridge, CB2 0QQ, UK
4. National Health Service Blood and Transplant (NHSBT), Cambridge Biomedical Campus, Cambridge, CB2 0PT, UK
5. BHF Centre of Excellence, Division of Cardiovascular Medicine, Addenbrooke's Hospital, Cambridge Biomedical Campus, Cambridge, CB2 0QQ, UK
6. Servicio de Hematología, Hospital Universitario de Salamanca, Salamanca, Spain.
7. Servicio de Hematología, Complexo Hospitalario Universitario de A Coruña, A Coruña, Spain
8. Laboratory of Haematology, La Timone Hospital, Marseille, France; C2VN, INRAE, INSERM, Aix-Marseille Université, Marseille, France
9. National Heart, Lung and Blood Institute's. The Framingham Heart Study, Framingham, MA, US
10. Institute of Cardiology, Jagiellonian University Medical College and John Paul II Hospital, 80 Prądnicka St, Kraków, Poland.
11. Banc de Sang i Teixits, Barcelona, Spain; Vall d'Hebron Research Institute, Universitat Autònoma de Barcelona (VHIR-UAB), Barcelona, Spain; CIBER de Enfermedades Cardiovasculares, Madrid, Spain
12. Department of Medical Genetics, University of Cambridge, Cambridge Biomedical Campus, Cambridge, UK
13. These authors contributed equally to this work.

##### Corresponding authors

### Methods

#### Genetic diagnostic methods

Genetic diagnostic methods used to evaluate *SERPINC1* gene included: i) PCR amplification and Sanger sequencing of exons and flanking regions, ii) Multiplex Ligation-dependent Probe Amplification (MLPA) covering the 7 exons of this gene, ii) whole gene sequencing by Ion Torrent technology (PGM; Thermo Fisher Scientific, Waltham, MA, USA) , iv) long-range PCR (LR-PCR) amplification of the whole gene followed by Next Generation Sequencing (NGS) MiSeq platform (Illumina, San Diego, CA, USA) and/or v) Comparative Genomic Hybridization array (CGHa; CytoScan® HD Array; Thermo Fisher Scientific). These methods were performed as previously described (1, 2).

#### Sequencing and basecalling

Long-read WGS was done using the PromethION platform (Oxford Nanopore Technologies). Samples were prepared using the 1D ligation library prep kit (SQK-LSK109), and genomic libraries were sequenced on R9 flow cell. Read sequences were extracted from base-called FAST5 files by Guppy (versions 3.0.4 to 3.2.8; 3.0.4+e7dbc23 to 3.2.8+bd67289) to generate FASTQ files, that were then merged per sample.

#### Data processing

We developed an *in-house* SV discovery workflow inspired on rules from nano-snakemake (3, 4), which is publicly available at https://github.com/who-blackbird/magpie. An overview of the workflow is shown in Figure 1A.

##### SV identification

Reads were aligned against the GRCh38/hg38 human reference genome using minimap2 (2.17-r941) with default parameters for nanopore data (`‘-ax map-ont’` parameter). SV discovery was done using a combination of three different algorithms:

- Sniffles v1.0.11 was executed with a supporting read evidence of 4 (‘-s 4’ parameter) due to coverage variability.
- NanoSV v1.2.4 was executed with default parameters. It was run on each independent chromosome in parallel to optimize compute time, with the limitation that inter-chromosomal variants were not detected.
- SVIM v1.2.0 was executed with default parameters. Resulting SV calls with a quality score of less than 10 were filtered out of the dataset and not used for analysis.

SV calls were merged at two different levels: intra-sample merge, to merge calls within individuals that had been identified by all three algorithms, and inter-sample merge, to merge SV calls across individuals.

Intra-sample merge. For each of the 19 samples, SV calls from all three different algorithms were merged using SURVIVOR v.1.0.7. VCF files were concatenated into one using bcftools (5), and SURVIVOR was run using the command '`SURVIVOR merge in.fofn 500 1 -1 -1 -1 -1 out.vcf`', requiring a maximum distance of 500bp between breakpoints. Additionally, intra-sample merge was done independently of the SV type, since different SV detection algorithms determine the type in different ways. For example, Sniffles determines canonical (DEL, DUP, INS, INV, TRA) and some complex SV types, while NanoSV calls only breakends (BND). The following options were turned-off: take the strands of SVs into account, estimate distance based on the size of SV, minimum size of SVs to be taken into account. After running SURVIVOR, an 'in-house' script was used to select the most common SV type. If there was no common type, the order of selection was NanoSV (if the SV type was not a BND) > Sniffles > SVIM type.

Inter-sample merge. Then, all the 19 samples were merged using SURVIVOR, taking the SV type into account. The command run was: '`SURVIVOR merge in.fofn 500 1 1 -1 -1 -1 out.vcf`'.

**Identification of candidate SVs in *SERPINC1***

After filtering out variants that overlapped bad mapping quality regions (obtained from (6)), a total number of 83,486 SVs were identified across all participants, with a median of 23,996 (sd $\pm$ 3,431) SVs per sample. In order to identify disease-causing SVs associated with antithrombin deficiency, we filtered for SVs overlapping the region [GRCh38/hg38] chr1:173,903,500-173,931,500, which includes *SERPINC1* gene and its promoter region. A total number of 99 SVs were observed, of which 10 were absent in gnomAD, NGC and the NIHR BioResource (6-8). These 10 SVs were observed in 9 samples: 6 were deletions, 1 was a tandem duplication, 1 was a complex SV formed by 1 duplication and 1 deletion, and 1 was a SINE-VNTR-Alu (SVA)-type retrotransposon insertion.

**Manual inspection of SVs**

Alignments for all the cases without a candidate variant were manually inspected at the above locus of interest using IGV (9). An additional SVA insertion was observed in P10 at the same position as the SVA insertion called in P9. In this sample, only two reads supported the alternate allele, explaining why the variant was not called in P10 by any of the variant callers used. Running Sniffles with a read evidence of 2 ('-s 2' parameter) on the P10 data resulted in the SVA insertion being called.

**Validations and breakpoint flanking sequence analysis**

All candidate SV junctions were confirmed by PCR amplification and Sanger sequencing (including the SVA insertion in P10 that had very low coverage) to resolve all variant configurations at nucleotide level resolution.

We then manually characterized the presence of microhomology, insertions and deletions at the breakpoints (Figure S4, Table S2) as previously described (10). The percentage of repetitive sequence was also calculated for each junction (+/− 150 bps) by intersecting these regions with the human genomic repeat library (hg38) from RepeatMasker version open-4.0.5 (http://www.repeatmasker.org) using bedtools (11) (Table S3).

***De novo* assembly**

*De novo* assembly was performed to characterize the SVA insertion in P9. Reads within the region [GRCh38/hg38] Chr1:173,840,000-174,820,000 were extracted from the alignment of this individual and converted to a FASTQ file using Samtools (5). *De novo* assembly was performed with wtdbg2 v2.5, using the parameters `` `-x ont -g 980k -X 10 -e 3` `` (12). The *de novo* contig was then aligned to the reference genome using minimap2 with default parameters for nanopore reads. The genomic sequence containing the SVA RE was then extracted from the alignment and analyzed with RepeatMasker (http://www.repeatmasker.org) to characterize the type of SVA and its subelements.

#### Supplementary figures

**Figure S1. Sequencing results colored by participant. (A)** Giga bases sequenced **(B)** Percentage of bases in genome sequenced at a specific minimum coverage **(C)** Median coverage in *SERPINC1* + promoter region **(D)** Coverage distribution in *SERPINC1* + promoter region **(E)** Percentage of reads with a minimum Q score **(F)** Read N50, which refers to a value where half of the data is contained within reads with alignable lengths greater than this.

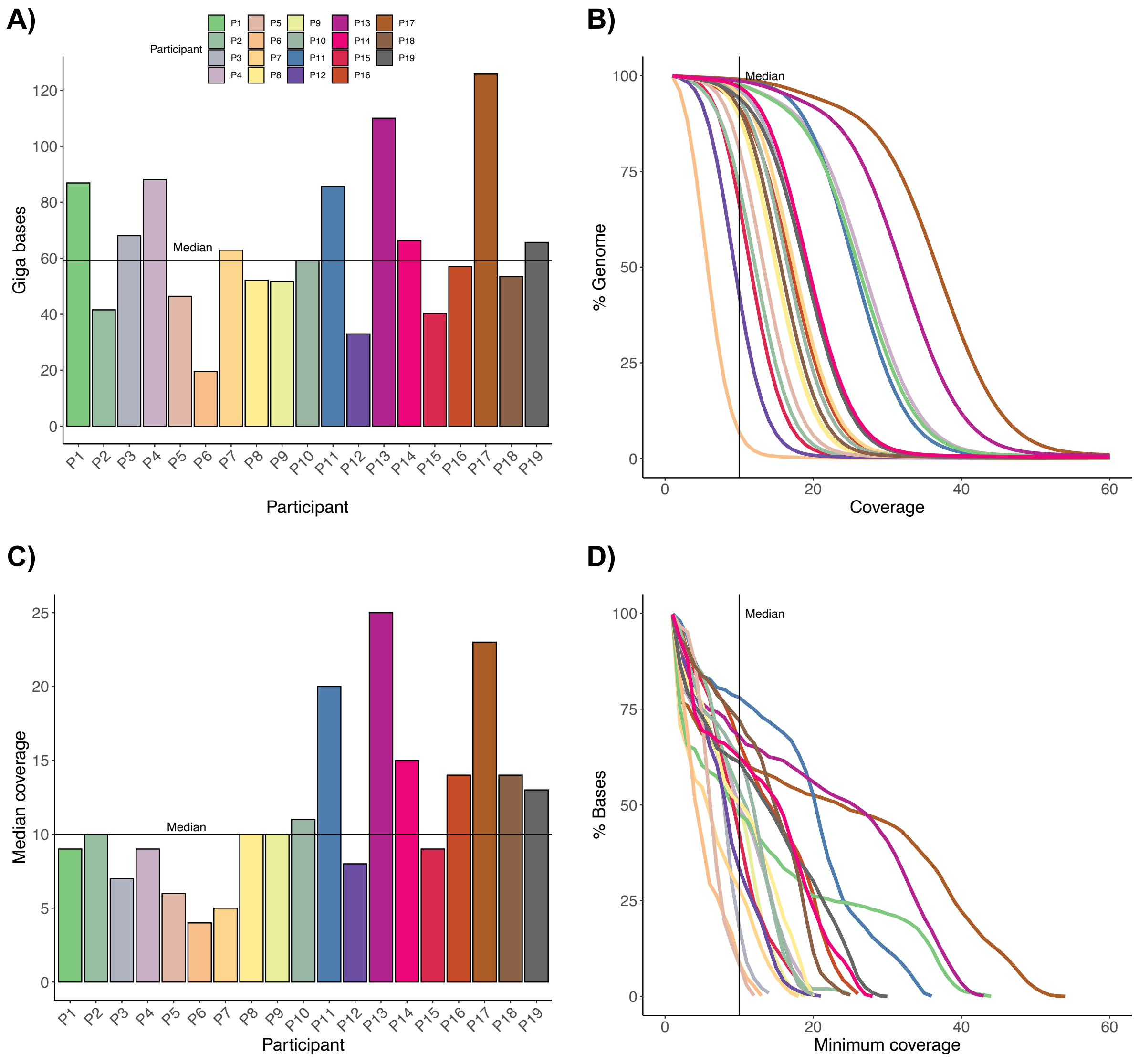

E)

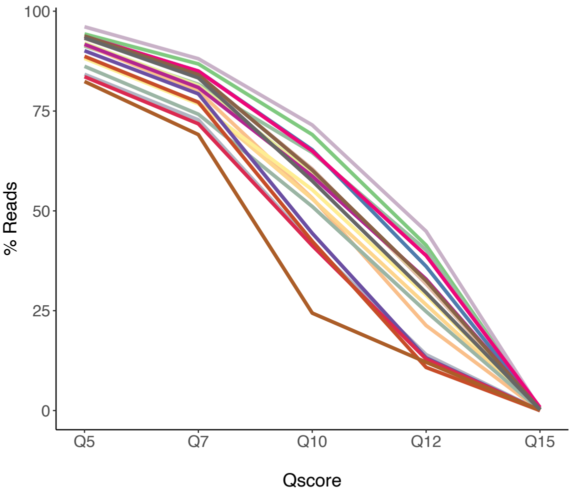
100
75
50
25
0
% Reads
Q5
Q7
Q10
Q12
Q15
Qscore

F)

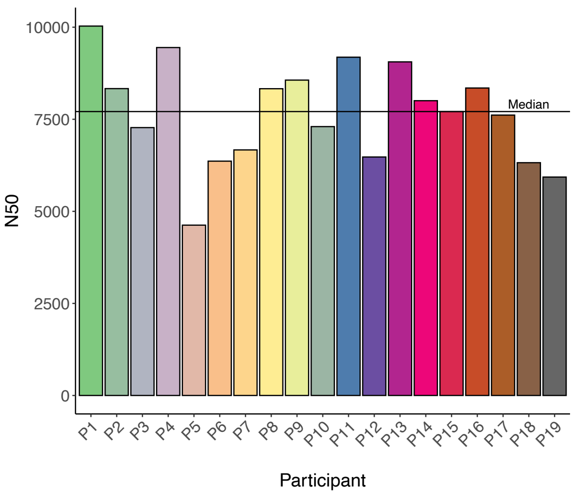
10000
7500
5000
2500
0
N50
Median
P1
P2
P3
P4
P5
P6
P7
P8
P9
P10
P11
P12
P13
P14
P15
P16
P17
P18
P19
Participant

**Figure S2. Structural variants metrics. (A)** Number of SVs identified by type and participant **(B)** Number of SVs by SV size **(C)** Fraction of SVs per allele count in our internal cohort of 62 individuals with long-read sequencing data **(D)** Number of SVs by median coverage and participant.

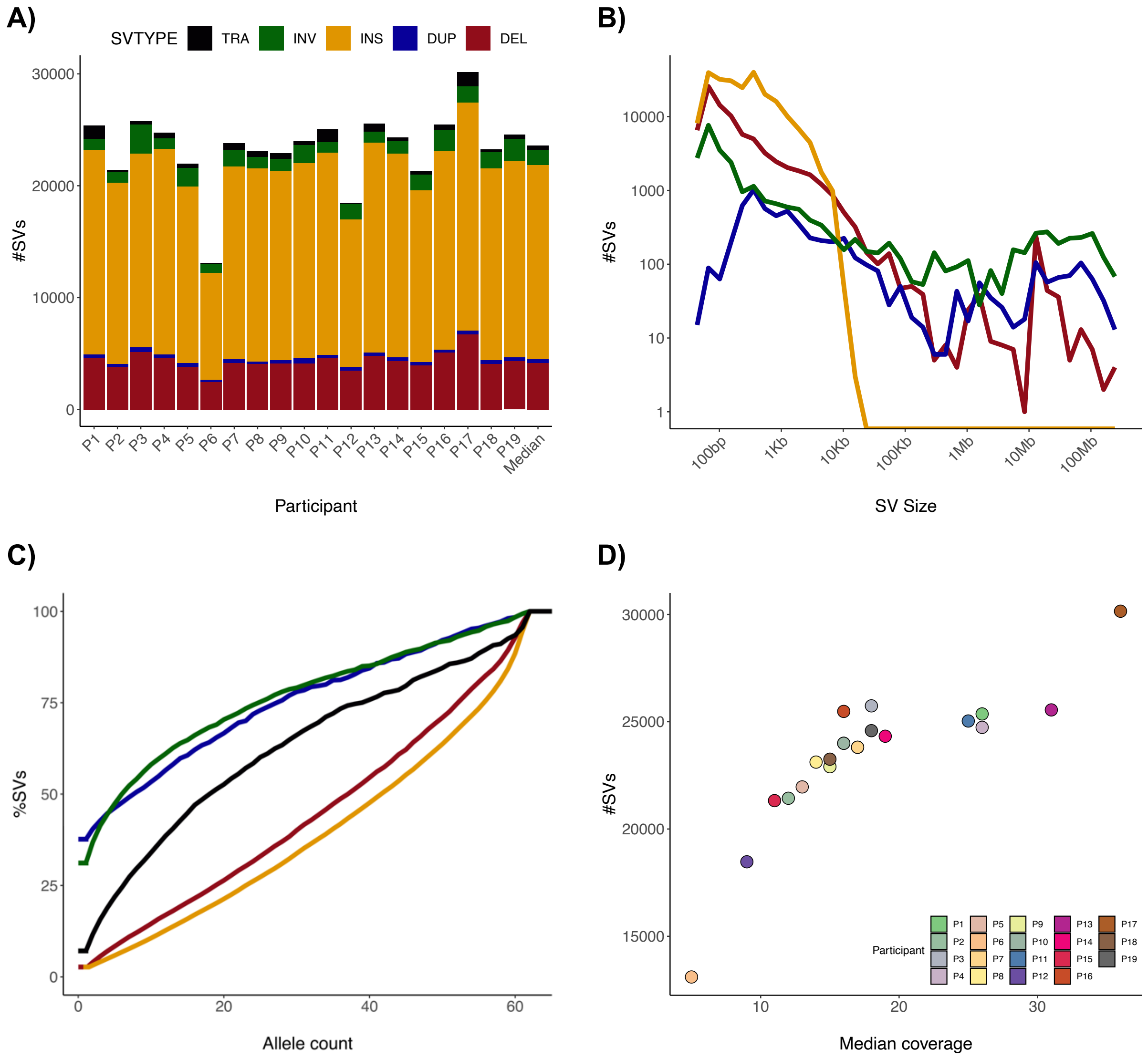

**Figure S3. SV resolution of P2 and P6.** Schematic representation of genetic diagnostic methods used to characterize the SVs in participants P2 **(A)** and P6 **(B).** Results from MLPA, LR-PCR and nanopore are shown in white boxes. Primers used for both LR-PCR and Sanger validation experiments are shown with orange and green arrows respectively. If present, J1 and J2 correspond to the new formed junctions described in Figure S6. J=New junction; M=Molecular weight marker; P=patient; C=control; B=Blank. **(B)** For the LR-PCR results, C1 and P1 correspond to PCR 1 (done with Primer F + Primer R), and C2 and P2 correspond to PCR2 (done with Primer F + Primer R2).

**A)**

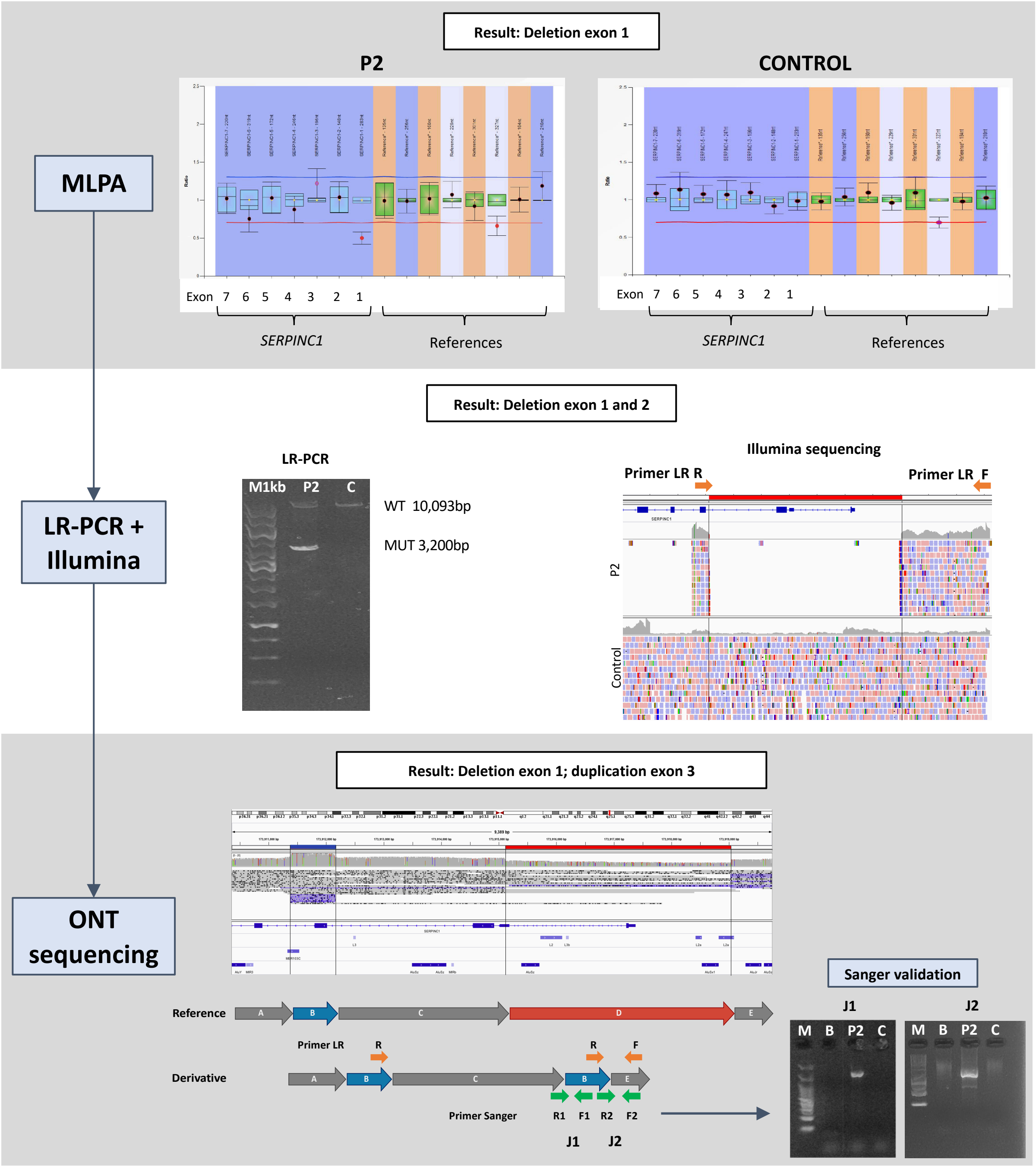

B)

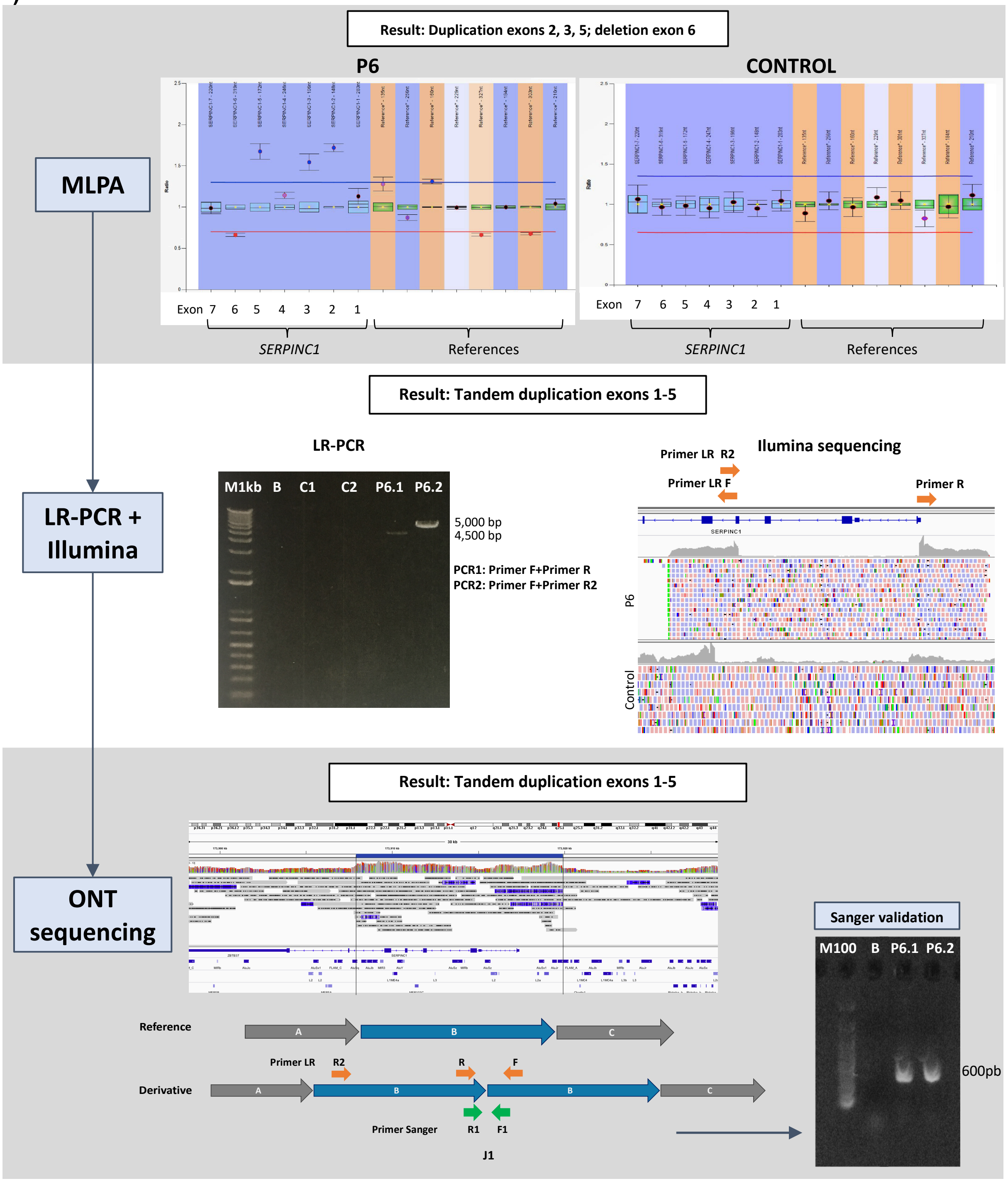
Result: Duplication exons 2, 3, 5; deletion exon 6
P6
CONTROL
MLPA
Exon 7 6 5 4 3 2 1
SERPINC1
References
Result: Tandem duplication exons 1-5
LR-PCR
M1kb B C1 C2 P6.1 P6.2
5,000 bp
4,500 bp
PCR1: Primer F+Primer R
PCR2: Primer F+Primer R2
Ilumina sequencing
Primer LR R2
Primer LR F
Primer R
SERPINC1
P6
Control
LR-PCR + Illumina
Result: Tandem duplication exons 1-5
ONT sequencing
Sanger validation
M100 B P6.1 P6.2
600pb
Reference
A
B
C
Derivative
Primer LR
R2
R
F
Primer Sanger
R1
F1
J1

**Figure S4**. **SVA sequence alignments. (A)** The consensus sequences of SVA-A, -B, -C, -D, -E, and -F were taken from RepeatMasker (http://www.repeatmasker.org), then aligned using MAFFT (13) with default parameters. SVA_query corresponds to the SVA insertion in P9. Alignments were visually inspected and colored by nucleotide using JalView (14). Sub-elements of the SVAs are indicated underneath the consensus sequence matching colors in Figure 2C. **(B)** A phylogenetic tree was constructed with the Neighbour-Joining (NJ) algorithm using the Jukes-Cantor substitution model and visualized with iTol (15). The SVA insertion in P9 was observed to be closest to the SVA E in the phylogenetic tree.

**(A)**

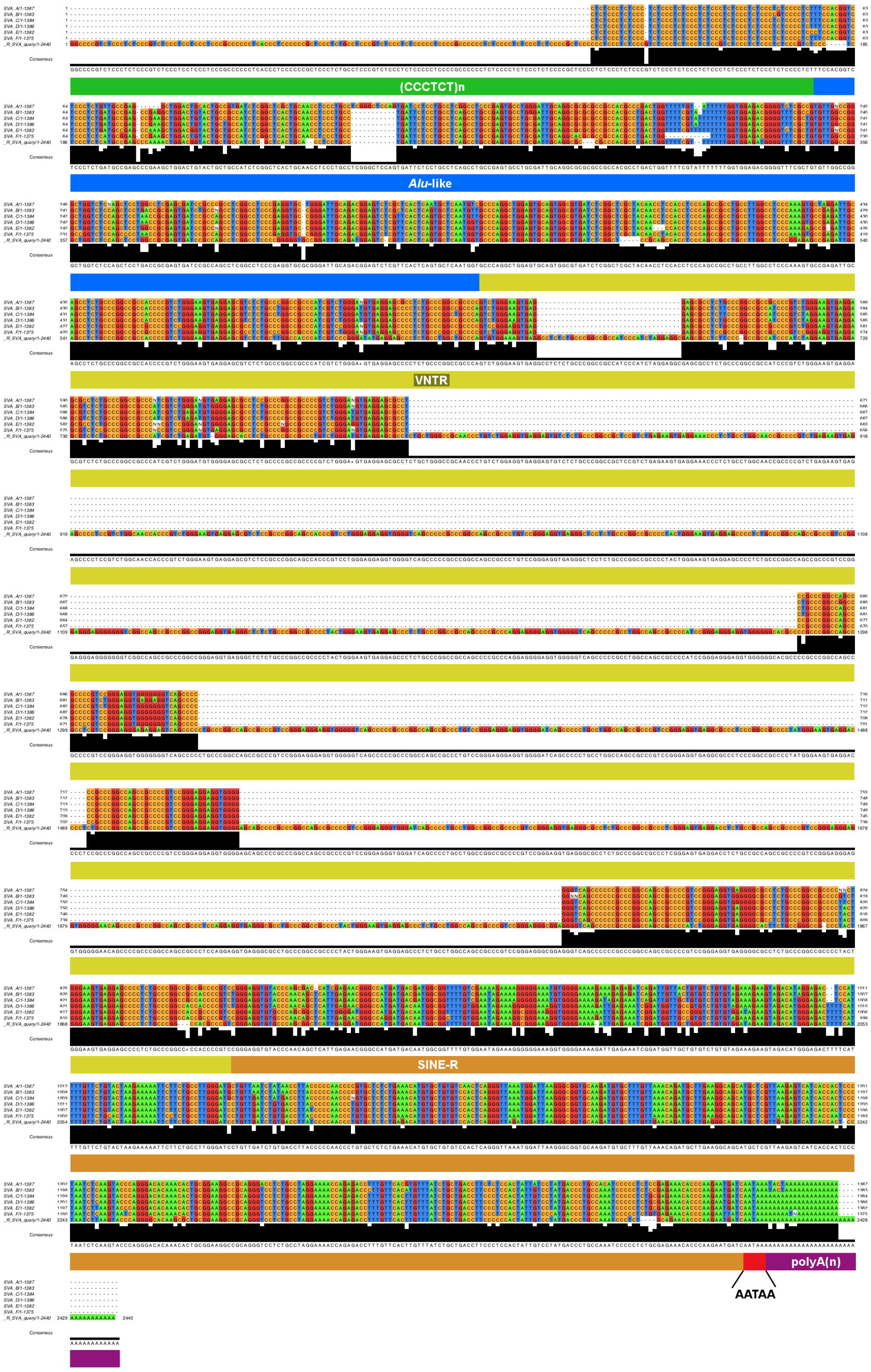

**(B)**

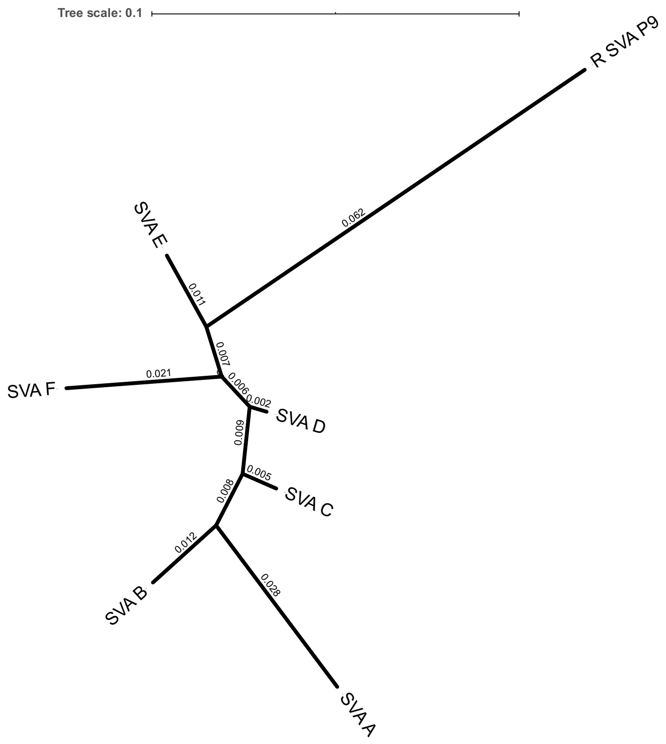
Tree scale: 0.1
R SVA P9
0.062
SVA E
0.011
0.007
SVA F
0.021
0.006
0.002
SVA D
0.009
0.005
SVA C
0.008
0.012
SVA B
0.028
SVA A

**Figure S5. PCR amplification validation of the SVA insertion in P9 and P10. (A)** Schematic of *SERPINC1* gene (NM_000488) with zoom to intron 6 showing the SVA structure. Primers used in the long- and short-range PCRs are shown in orange and green respectively. Primer 8* was specifically designed within the inserted SVA sequence. Briefly, four reads of the retrotransposon present in the nanopore data for P9 and P10 were aligned to identify regions without any mismatch in order to select a 20 nucleotide sequence to be used as primer. That sequence was also checked to be present in *de novo* assembly alignment. **(B)** Primer combinations for PCR amplifications and expected sizes for wild type and mutated alleles are shown in the table. PCRs 1-4 were tested under different experimental conditions, and although in all cases the wild type allele was always amplified, no amplification of the mutated allele containing the SVA was obtained in P9 or P10. **(C)** The amplification of PCR 4 in agarose gel is shown. Only the 800 base pairs (bp) of the wild type allele was amplified in P9, P10 but also a healthy control. Only PCR 5, using the primer specific of the SVA rendered positive results and a specific 550bp band was obtained in P9, P10 and two relatives. **(D)** Family pedigrees of P9 and P10, including clinical information, the diagnosis of antithrombin deficiency (semi-filled symbols) and the anti-FX activity (as % of a reference plasma). B=Blank; M=Molecular Weight Marker; DVT=Deep vein thrombosis; PE=Pulmonary embolism.

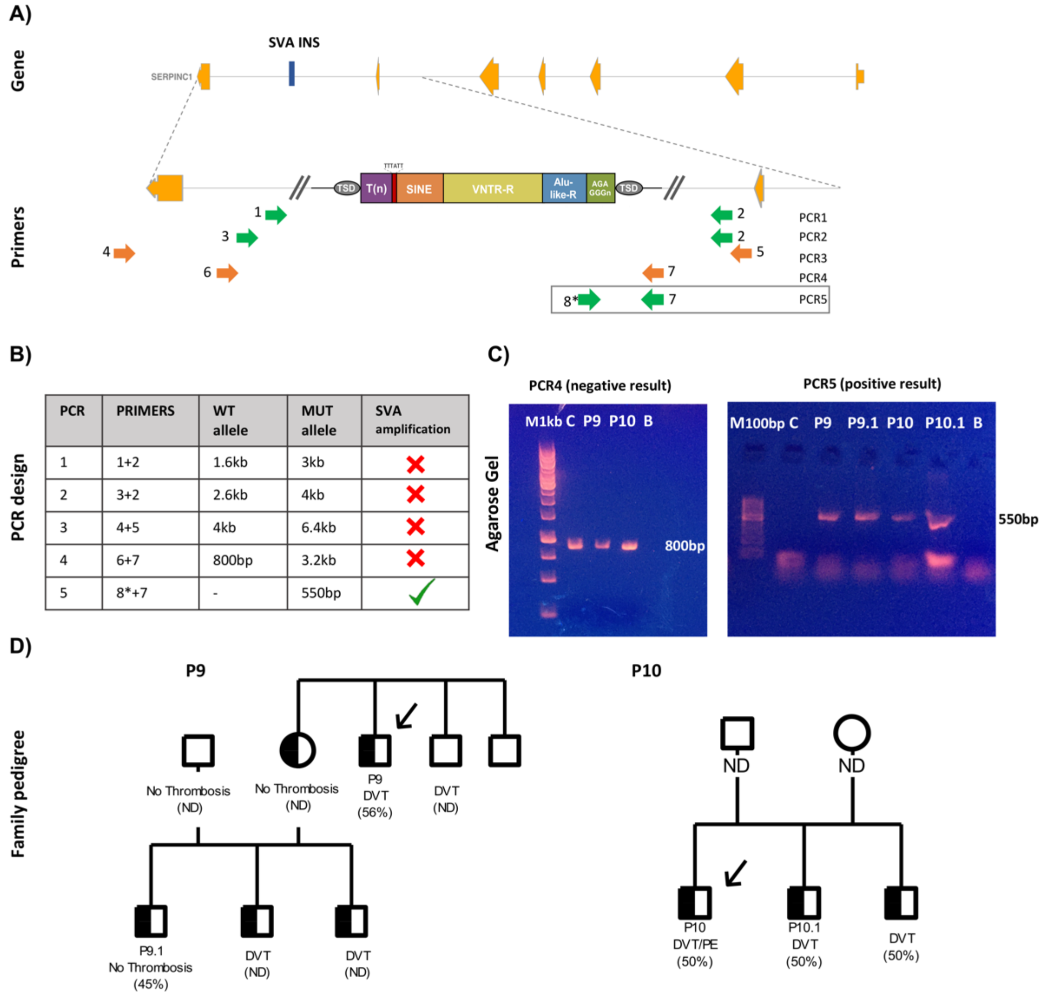

| PCR | PRIMERS | WT allele | MUT allele | SVA amplification |
|---|---|---|---|---|
| 1 | 1+2 | 1.6kb | 3kb | ✗ |
| 2 | 3+2 | 2.6kb | 4kb | ✗ |
| 3 | 4+5 | 4kb | 6.4kb | ✗ |
| 4 | 6+7 | 800bp | 3.2kb | ✗ |
| 5 | 8*+7 | - | 550bp | ✓ |

**Figure S6. Nucleotide level characterization of the candidate SVs.** Breakpoint junction sequence is aligned to the proximal and distal genomic reference sequence. Alignment is only shown for novel breakpoint junctions in the derivative chromosome. Microhomology at the breakpoint is indicated in red. Sequence in blue indicates inserted sequences at the breakpoint junction. Underline indicates repetitive elements in the reference, specified in Italic. J=Junction. **(A-H)** P1-8, **(I)** P9 and P10.

## (A) P1

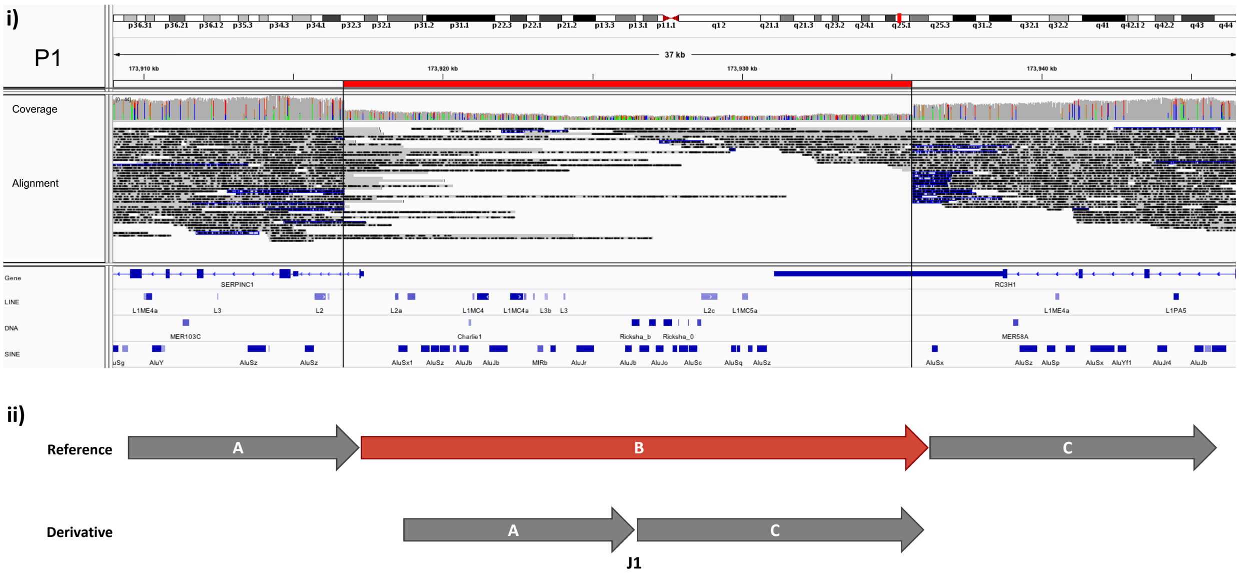

**iii)**

```
1-173916654   CCAAGACAGTTGTGCCACCACCCCGTTTCTAGCCTCCTGTGACAACTCA*GATGAAGAAGGGAGTGTGTGTGGCATTGAGGGAGGGACAACTCTCATTGTG   173916754
J1-1          CCAAGACAGTTGTGCCACCACCCCGTTTCTAGCCTCCTGTGACAACTCA*GGAGCAAGGAACAAGGAAAAAAAGCAAAAAACAAAAAACAAATTTTGGGGG   100
1-173935659   AAATATGCCATTTGTCCTTACAGGCTCATGATATCACAGAGAAAAGGAA*------------AAGGAAAAAAAGCAAAAAACAAAAAACAAATTTTGGGGG   173935759
```

*A-rich*

## (B) P2

i)

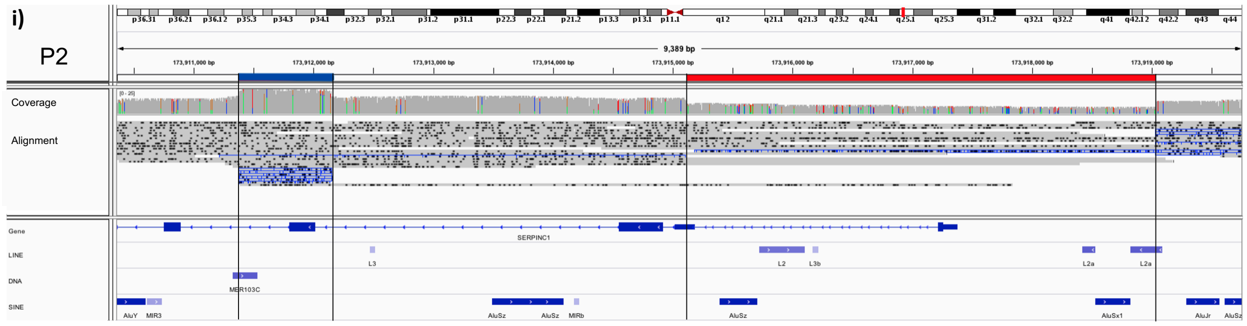

ii)

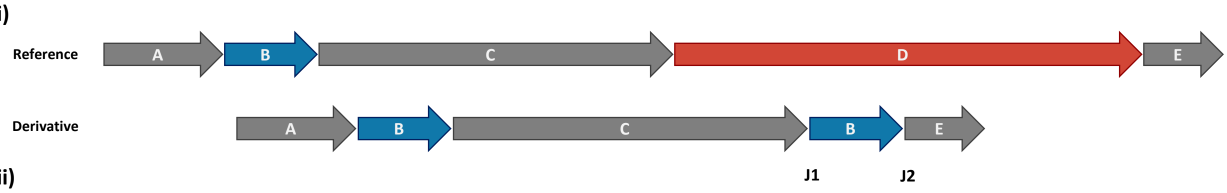

iii)

```
1-173915066   CCGGGACAGGTTCAGTCCTAGACTTCTTGCCAGGGGACAGTTCAGTTGC*CTGGACGTGGTCATGTTGGAATTAGAATCATCTGATGGTTATTTGGAGTA   173915165
J1-1          CCGGGACAGGTTCAGTCCTAGACTTCTTGCCAGGGGACAGTTCAGTTGC*-----AGAGTTTCTCAGAACCTTCATAGGCCCATGTGCTTCATGAATCTC   94
1-173911329   TCAAATACATTTGAGGAATGCCGATTGAATAAAGCCAACAGATTTCTTT*GTTGCAGAGTTTCTCAGAACCTTCATAGGCCCATGTGCTTCATGAATCTC   173911428
                                  MER103C                                            MER103C

1-173912101   AATAGCCACCTCAGTTGTTAACTCCTTCAAGCACAGTACCTGGGACAGC*ATAAATGCTCAACTCTTGTGTTCTGATGAATAAAAGGATAAAAGAAGAATG   173912201
J2-1          AATAGCCACCTCAGTTGTTAACTCCTTCAAGCACAGTACCTGGGACAGC*---TGTCCAAAGAGAAATCAGAAATACAAAGACCTTGAGGTTTCAGGAACA   97
1-173918984   GAAACCAGAATGAAATGATGGAGGGGGCCATATGAATATCTTTAGGAAG*AGCTGTCCAAAGAGAAATCAGAAATACAAAGACCTTGAGGTTTCAGGAACA   173919084
                                  L2a                                                L2a
```

## (C) P3

i)

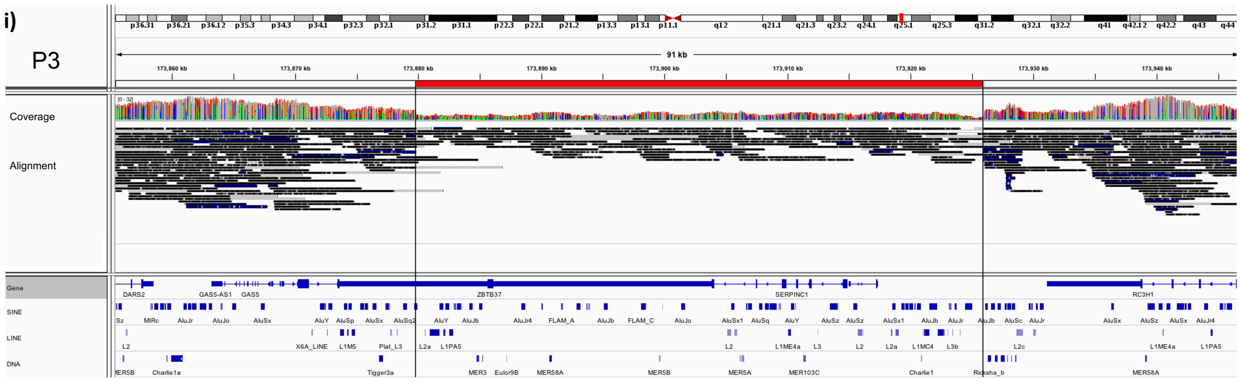

ii)

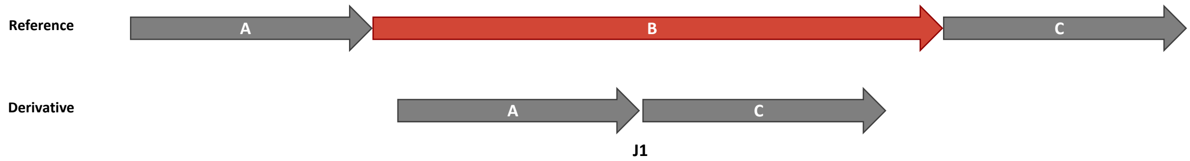

iii)

```
                          AluSx1                                                AluSx1
1-173879770   GTGAAACCCCATCTCTACTAAATATAAAACATTAGCCAGGCATGGTGGT*GGGCACCTGTAATCTCAGCTACTCGGGAGGCTGCAACAGGAGAATCACTTG   173879870
J1-1          GTGAAACCCCATCTCTACTAAATATAAAACATTAGCCAGGCATGGTGGT*GCCTGGACAAGGCTATTCTTTGGAGGCTGAGCAAGAGAGAGTTGTGTGGGG   100
1-173925939   AAATGCTTACTCATCCAGAAGAATTGACACCTTGTATGAACAGAAACAT*---TGGACAAGGCTATTCTTTGGAGGCTGAGCAAGAGAGAGTTGTGTGGGG   173926039
```

## (D) P4

i)

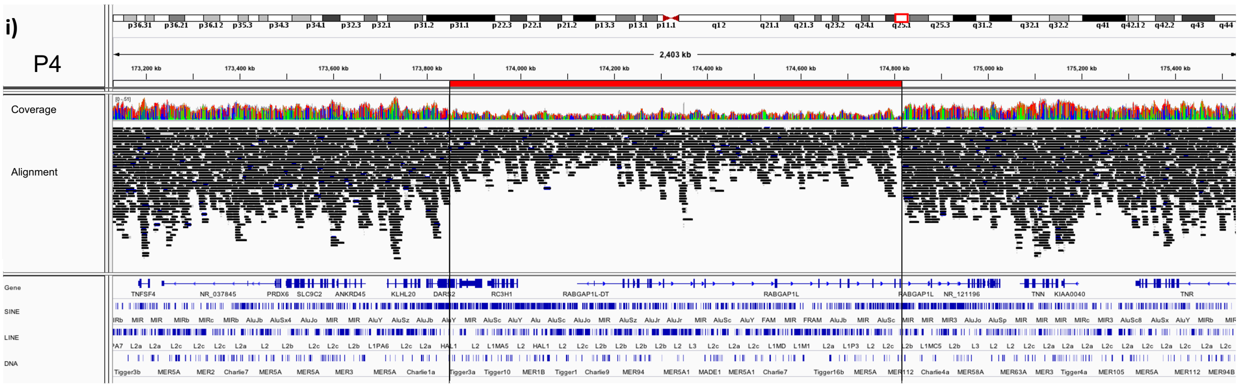

ii)

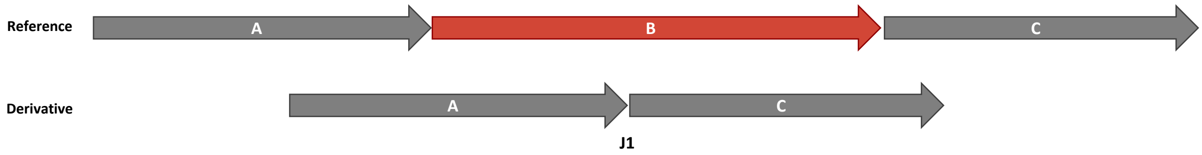

iii)

```
                HAL1                                                     AluY
1-173847797  CCTCACCTAGTTACTCTGTACCTTCTGCACAGCTGTCTTTCTGAACCTTT*TTTTTTTTTTTTTTTTTTTTTTTGAGACGGGAGTCTCACTCTGTCGCCCAG  173847897
J1-1         CCTCACCTAGTTACTCTGTACCTTCTGCACAGCTGTCTTTCTGAACC---*TTTTTTTTTTTTTTTTTTTTTTTGAGATAGAGTTTTGCTCTTGTTGCCCAG  97
1-174816097  TTGTATGAACATATGAGCATTTTGTTTTTCTTAGTAACATAATACATAGC*TTTTTTTTTTTTTTTTTTTTTTTGAGATAGAGTTTTGCTCTTGTTGCCCAG  174816197
                                                                         AluSp
```

## (E) P5

i)

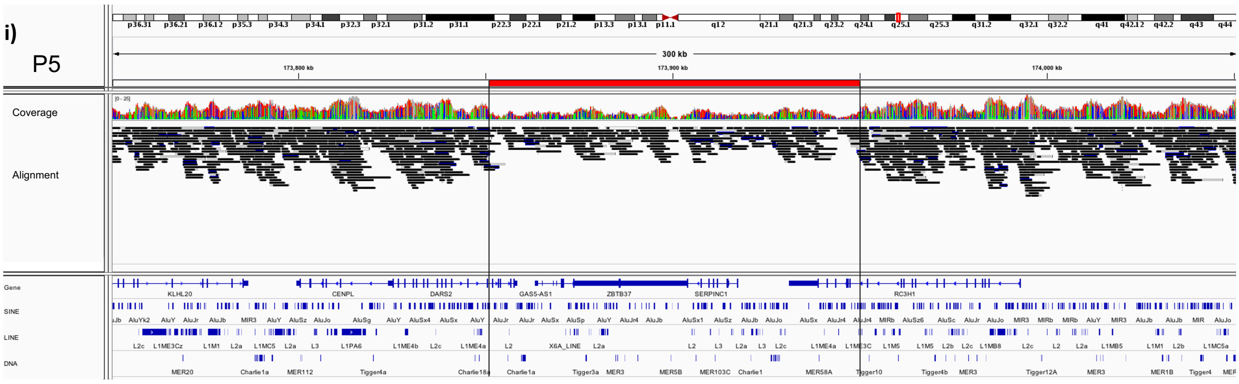

ii)

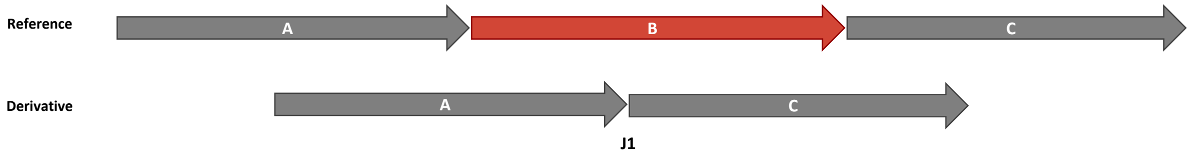

iii)

```
                AluSx1                                                   AluSx1
1-173850946  TTAAGGGTTGGCCGGGCATGGTGGCTAACGCCTGTAATCCCAGCACTTT*GGGAGGCCGAGGTGGGTGGATCGCCTGAGGTCAGGAGTTTGAGACCAGCCT  173851046
J1-1         TTAAGGGTTGGCCGGGCATGGTGGCTAACGCCTGTAATCCCAGCACTTT*-AAGTACACATCTTAAAAGCAATGCTAAGTGAAAACATGAAAATGCAAGCT  99
1-173950124  TGCCACTGCACTCCAGCCTGGGTACAGAGTGGAACTCCGTCTCAAAAAA*AAAGTACACATCTTAAAAGCAATGCTAAGTGAAAACATGAAAATGCAAGCT  173950224
                AluSx                                                    L1ME3C
```

## (F) P6

i)

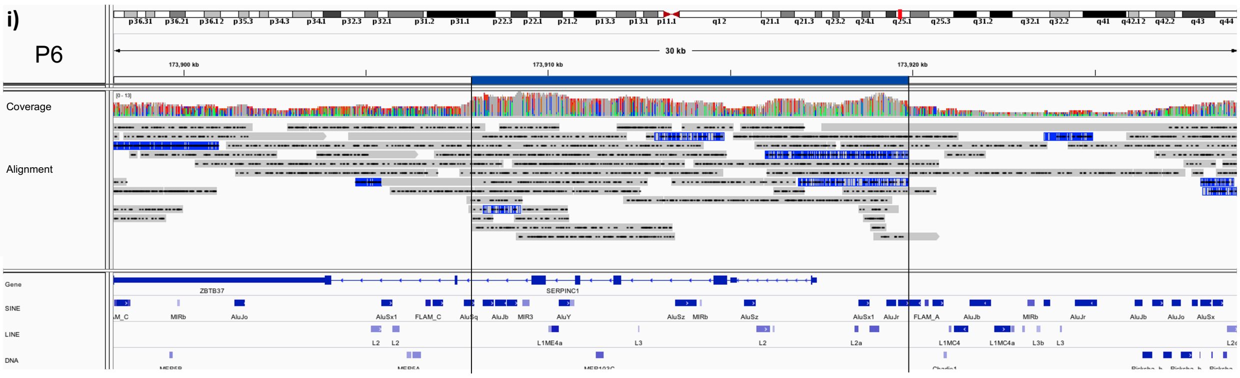

ii)

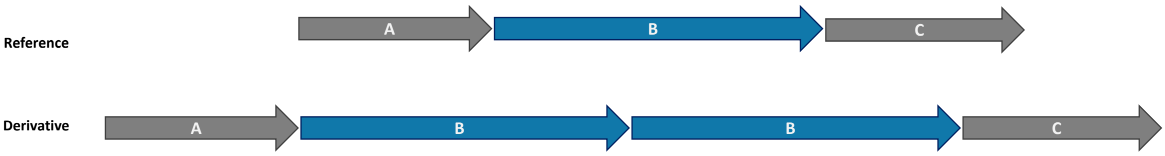

iii)

```
                 AluSz                                         AluSz
1-173919740   ACCTGTAATCTCAGCTACTCAGGAGGCTGAGGCAGGAGAATCACTTGAA*CCCGGGAGGTGGAGGCTGCAGTGAGTCGAGATTACGCCACTGCACTCTAGC   173919840
J1-1          ACCTGTAATCTCAGCTACTCAGGAGGCTGAGGCAGGAGAATCACTTGAA*TCTGGGGGGTGGAGGTTGCAGTGAGCCAAGATTGCACCACTGCACTCCAGC   100
1-173908362   GCCTGTAATCCTAGCTCCTTGGGAGGCTGAGGCAGGAGAATCACTTGAA*TCTGGGGGGTGGAGGTTGCAGTGAGCCAAGATTGCACCACTGCACTCCAGC   173908462
                 AluSx1                                        AluSx1
```

## (G) P7

i)

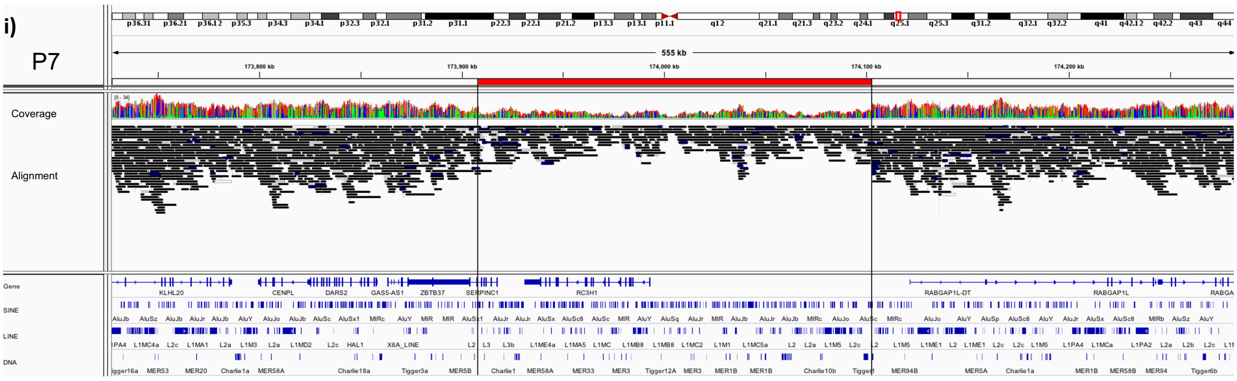

ii)

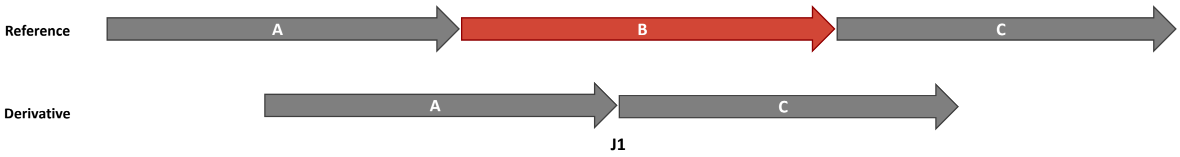

iii)

```
                 AluSx1                                        AluSx1
1-173908284   TTCAAGACCAGCCTGGCCAACATAGTGAAACCCCGTTTCTACTAAAAAT*GCAAAAATTAGCCGGGCGTGGTGGCAGGCGCCTGTAATCCTAGCTCCTTGG   173908384
J1-1          TTCAAGACCAGCCTGGCCAACATAGTGAAACCCCGTTTCTACTAAAAAT*ACAAAAATTAGCCAGGCATGGTGGTGAGCGCCTGTAATCCCAGCTACTCCA   100
1-174102965   GTCAAGACCAGCCTGATCAACATGGAGAAACACCGTCTCTACTAAAAAT*ACAAAAATTAGCCAGGCATGGTGGTGAGCGCCTGTAATCCCAGCTACTCCA   174103065
```

## (H) P8

i)

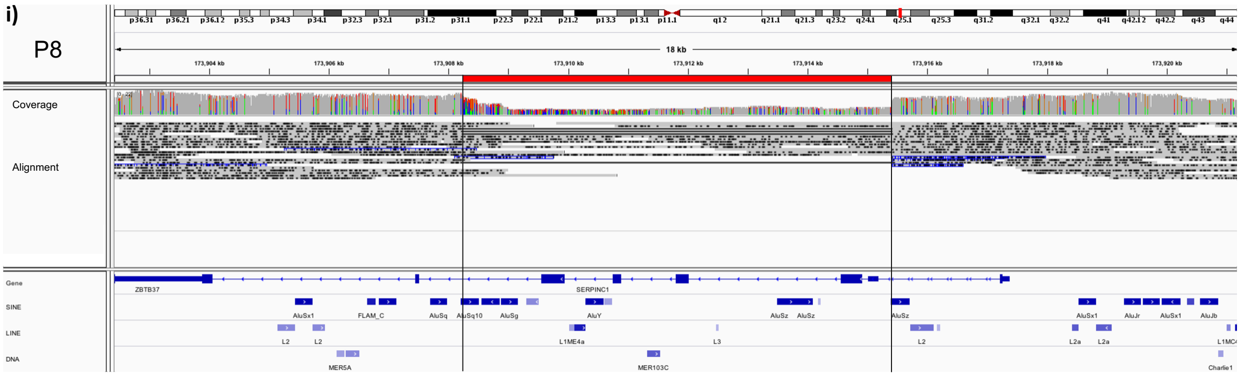

ii)

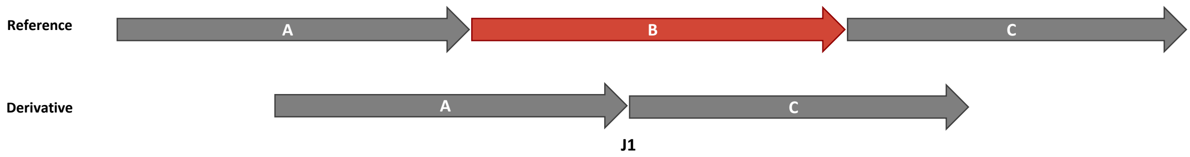

iii)

```
                                                                   AluSx1
1-173908168   GGCAACATTTTTAGTTTTCAAAACTAGCTTAAAGACAGCCCAATAGCAC*AGTGGCTCACGCCTGTAATCCCAGCACTTTGGGAGGCCAAGGCAGGTAGAT   173908268
J1-1          GGCAACATTTTTAGTTTTCAAA-CTAGCTTAAAGACAGCCCAATAGCAC*AGTGGCTCACGCCTGTAATCCCAGCACTTTGGGAGGCTGAGGCATGCAGAT   99
1-173915355   AGAAGTAAGAAGAAATGGATGGAGCAAAGGCAATATCTGGCCGGGTGCG*-GTGGCTCACGCCTGTAATCCCAGCACTTTGGGAGGCTGAGGCATGCAGAT   173915454
                                                                   AluSp
```

#### (I) P9 and P10

i)

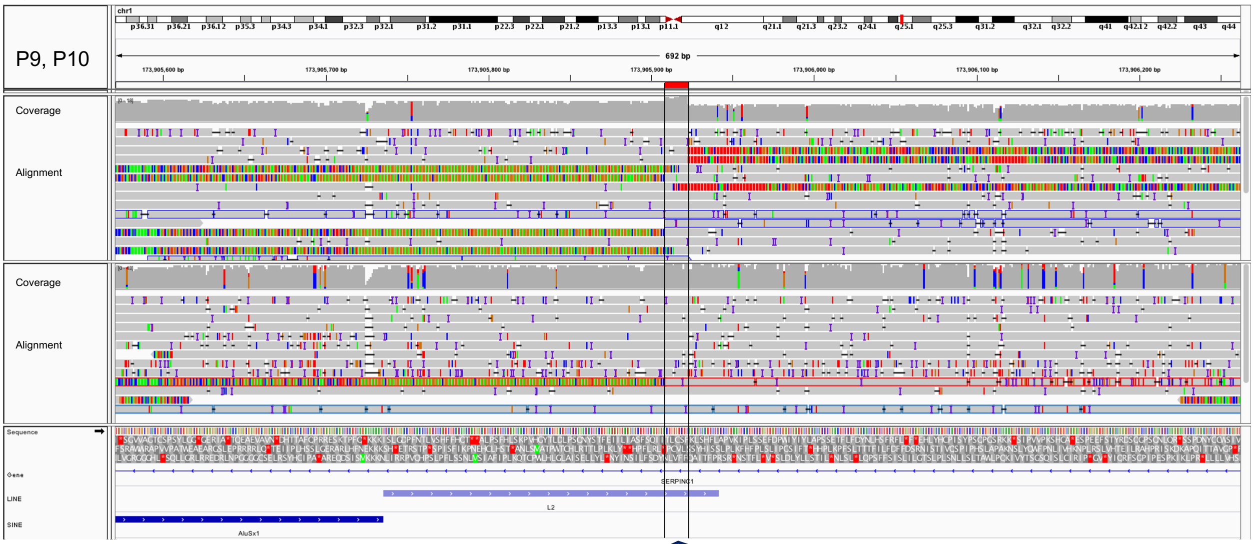

ii)

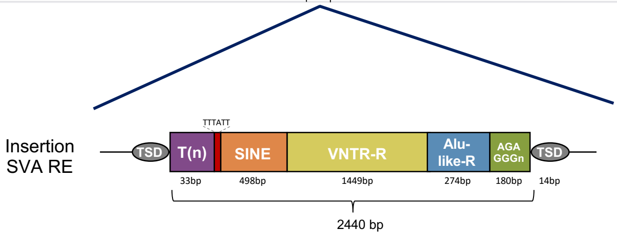

**iii)**

**Participant 9**

**P9 39-52:** **TSD**
P9 53-84: (T)n
P9 85-90: TTTATT sequence
P9 91-589: SINE
P9 590-2038: VNTR-R
P9 2039-2312: Alu-like-R
P9 2313-2492: (AGAGGG)n
**P9 2493-2506:** **TSD**

```
1-173905870    TTGAAATTATATTAATAGCATCCTTTTCTCAGATTATAACCTTGTGTTCTTT------------------------------------------------    173905926
               ||||||||||||||||||||||||||||||||||||||||||||||||||||
P9-1           TTGAAATTATATTAATAGCATCCTTTTCTCAGATTATAACCTTGTGTTCTTTTTTTTTTTTTTTTTTTTTTTTTTTTTTTTTTTTTTTATTGATCATTCT    100

1-173905926    ----------------------------------------------------------------------------------------------------    173905926

P9-101         TGGGTGTTCTGCAGAGGGATTTGGCAGGGTCATAGGACAATAGTGGGGGGAAGGTCAGCAGATAAACAAGTGAACAAAGGTCTCTGGTTTTCCTAGGCAG    200

1-173905926    ----------------------------------------------------------------------------------------------------    173905926

P9-201         AGGACCCTGCGGCCTTCCGCAGCGCTTGTGCCCCTGGGTACTTAAGATTAGGAGTGGTGATGACTCTCAACGAGCATGCTGCCCTCAAGCATCTGTTCAA    300

1-173905926    ----------------------------------------------------------------------------------------------------    173905926

P9-301         CAAAGCACATCTTGCACCGCCCTTAATCCATCTAACCCTGAGTGGACACAGCACATGTCTCAGAGAGCACAGGGTTGGGGATAAGGTCACAGATCAACAG    400

1-173905926    ----------------------------------------------------------------------------------------------------    173905926

P9-401         GATCCCAAGGCAGAAGAATTTTTCTTAGTACAGAACAAAATGAAAAGTCTCCCATGTCTACTTCTATCCACACAGACCCAGCAACCATCCGATTTCTCAA    500

1-173905926    ----------------------------------------------------------------------------------------------------    173905926

P9-501         TTTTTCCCCACCCTTCCCGCCTTTCTATTCCACAAAACCGCCATTGTCATCATGGCCCATCCTCAATGAGCCGCTGGGCACACCTCCCGACGGGCGTGGC    600

1-173905926    ----------------------------------------------------------------------------------------------------    173905926

P9-601         CGGGCAGAGGGGCTCCTCACTTCCCAGTAGGGCGGCCCGGCAGAAGTGCCCCTCACCTCCCAGATGGGGCGGCTGGCCGGGCGGGGGCTGACCCCTCCGC    700

1-173905926    ----------------------------------------------------------------------------------------------------    173905926

P9-701         CCTCCCGGACGGGCGGCTGGCCAGGCAGAGGGCTCCTCACTTCCCAGTAGGGGCGGCCGGGCAGGCGCCCTCACCTCCTGGAGGGCGGCTGGCCGGGCGG    800

1-173905926    ----------------------------------------------------------------------------------------------------    173905926

P9-801         GGCTGTTCCCCCACCTCCCTCCCGGACGGGGCGGCTGGCGGCAGAGGTCCTCACTCCCGAGGGCGGCCGGGCAGAGGCGCCCTCACCTCCCGGACGGGGC    900

1-173905926    ----------------------------------------------------------------------------------------------------    173905926

P9-901         GGCCGGCCAGGCAGGGGCTGATCCCACCCTCCCGGACGGGGCGGCTGGCCGGGCGGGGCTGCTCCCCACCTCCTCCCGGACGGGGCGGCTGGCCGGGCAG    1000

1-173905926    ----------------------------------------------------------------------------------------------------    173905926

P9-1001        AGGGGTCCTCACTTCCCATAGGGGCGGCCGGGAGGGCGCCTCACCTCCCGGACGGGCGGCTGGCCAGGCAGGGGGCTGATCCCCACCTCCCTCCCGGACA    1100

1-173905926    ----------------------------------------------------------------------------------------------------    173905926

P9-1101        GGGCGGCTGGCCGGGCGGGGGCTGACCCCCACCTCCCTCCCGGACGGGCGGCTGGCCGGGCAGGGGGCTGACTCCTCTCCCTCCCGGACGAGGCGGCTGG    1200

1-173905926    ----------------------------------------------------------------------------------------------------    173905926

P9-1201        CCGGGCGGGGCGTGCCCCCCACCTCCCTCCCGGATGGGGCGGCTGGCCAGGCGGGGGCTGACCCCCACCTCCCCTCCTGGGCGGGGCTGGCGGCCGGGCA    1300

1-173905926    ----------------------------------------------------------------------------------------------------    173905926

P9-1301        GAGGGCTCCTCACTTCCCAGTAGGGGCGGCCGGGCAGAGAGCCCTCACCTCCCGGCCGGGCGGCTGGCCGACCCCCCCTCCCTCCCGGACGGGCGGCTGG    1400

1-173905926    ----------------------------------------------------------------------------------------------------    173905926

P9-1401        CCGGGCAGAGGGGCTCCTCACTTCCCAGTAGGGGCGGCCGGGCAGAGGAGCCCTCACCTCCCGGACAGGGCGGCTGGCCGGGCGGGGGCTGACCCCACCT    1500

1-173905926    ----------------------------------------------------------------------------------------------------    173905926

P9-1501        CCTCCCAGGACGGGTGGCTGCCGGGCGGAGACGCTCCTCACTTCCCAGACGGGTGGTTGCCAGACGGAGGGGCTCTCACTTCTCAGACGGGGCGGTTGCC    1600

1-173905926    ----------------------------------------------------------------------------------------------------    173905926

P9-1601        AGGCAGAGGGTTTCCTCACTTCTCAGACGGAGCGGCCGGGCAGAGACACTCCTCACCTCCAGACAGGGTTGCGGCCCAGCAGAGGCGCTCCTCACATCCC    1700

1-173905926    ----------------------------------------------------------------------------------------------------    173905926
P9-1701        AGACAGGGCGGCGGGGCAGAGGTGCTCCCACATCTCAGACGATGGGCGGCCGGGCAGAGACGCTCCTCACTTCCTAGATGGGATGGCGGCGGGAAGAGGC    1800

1-173905926    ----------------------------------------------------------------------------------------------------    173905926

P9-1801        GCTCGCCTCCTAGATGGGATGGCGGCCGGGCAGAGAGGCCTCACTTTCCACACTGGGCAGCCAGGCAGAGGGCTCCTCATATCCCGGACGATGGGTGGC    1900
```

1-173905926 ---------------------------------------------------------------------------------------------------- 173905926

P9-1901 CAAGCAGAGACGCTCCTCACTTCCCAGACGGGGTGGCGGCCGGGCAGAGGCTGCAATCTCGGCTCTCCGGGAGGCCAAGGCAGGCGGCTGGGAGGTGGCT 2000

1-173905926 ---------------------------------------------------------------------------------------------------- 173905926

P9-2001 GCGGAGCCGAGATCACGCCACTGCACTCCAGCCTGGGCACCATTGAGCACTGAGTGAACGGACTCCATCTGCAATCCGGCACCCCCGGGGAGGCCGAGGC 2100

1-173905926 ---------------------------------------------------------------------------------------------------- 173905926

P9-2101 TGGCGGATCACTCGCGGCCAGGAGCTGGAGACCAGCCCGGCCAACACAGCGAAACCCCATCTCCACCAAAAAAACGAAAACCAGTCAGGCGTGGGCGGCG 2200

1-173905926 ---------------------------------------------------------------------------------------------------- 173905926

P9-2201 CCTGCAATCGCAGGCACTCGGCAGGCTGAGGCAGGAGAATCAGGCAGGAGGTGCAGTGAGCGAGATGGCAGCAGTACCGTCCAGTTTTGGGCTCGGCATG 2300

1-173905926 ---------------------------------------------------------------------------------------------------- 173905926

P9-2301 AGAGGGAGAGGGAGACGGGAGAGGGAGAGGGAGACGGAGAGGGAGAGGGAGACGGGAGAGGGAGGGGGGAGCGGGGAGAGGGAGAGGGGAGAGGGGGGCG 2400

1-173905926 ---------------------------------------------------------------------------------------------------- 173905926

P9-2401 GGGAGGGGGAGAGGGAGACGGGAGGCAGAGGGAGCGGGGGGAGGGTGAGGGGGGCGGGAGGGAGGAGGGAGACGGGAGAGGGAGACGGGGCC**ACCTTGTG** 2500

```
1-173905926   ------CAAGCTATCACATTTCCTCGCTCCCGTTAAAATTCCACTTTCCTCTGA   173905970
                    |||||||||||||| |||||  |||||||||||| |||||||||||||
P9-2501       TTCTTTCAAGCTATCACTTTTCCCTGCTCCCGTTAAA-TTCCACTTTCCTCTGA   2600
```
